# Self-Assembled Origami Neural Probes for Scalable, Multifunctional, Three-Dimensional Neural Interface

**DOI:** 10.1101/2024.04.25.591141

**Authors:** Dongxiao Yan, Jose Roberto Lopez Ruiz, Meng-Lin Hsieh, Daeho Jeong, Mihály Vöröslakos, Vittorino Lanzio, Elisa V. Warner, Eunah Ko, Yi Tian, Paras R. Patel, Hatem ElBidweihy, Connor S. Smith, Jae-Hyun Lee, Jinwoo Cheon, György Buzsáki, Euisik Yoon

## Abstract

Flexible intracortical neural probes have drawn attention for their enhanced longevity in high-resolution neural recordings due to reduced tissue reaction. However, the conventional monolithic fabrication approach has met significant challenges in: (i) scaling the number of recording sites for electrophysiology; (ii) integrating of other physiological sensing and modulation; and (iii) configuring into three-dimensional (3D) shapes for multi-sided electrode arrays. We report an innovative self-assembly technology that allows for implementing flexible origami neural probes as an effective alternative to overcome these challenges. By using magnetic-field-assisted hybrid self-assembly, multiple probes with various modalities can be stacked on top of each other with precise alignment. Using this approach, we demonstrated a multifunctional device with scalable high-density recording sites, dopamine sensors and a temperature sensor integrated on a single flexible probe. Simultaneous large-scale, high-spatial-resolution electrophysiology was demonstrated along with local temperature sensing and dopamine concentration monitoring. A high-density 3D origami probe was assembled by wrapping planar probes around a thin fiber in a diameter of 80∼105 μm using optimal foldable design and capillary force. Directional optogenetic modulation could be achieved with illumination from the neuron-sized micro-LEDs (μLEDs) integrated on the surface of 3D origami probes. We could identify angular heterogeneous single-unit signals and neural connectivity 360° surrounding the probe. The probe longevity was validated by chronic recordings of 64-channel stacked probes in behaving mice for up to 140 days. With the modular, customizable assembly technologies presented, we demonstrated a novel and highly flexible solution to accommodate multifunctional integration, channel scaling, and 3D array configuration.

## 1. Introduction

Enhancement of recording capability and integration of multi-modalities are two essential needs in neural probe development. High-channel-count neural probes have demonstrated their extraordinary practicality for high spatial resolution and large area coverage, allowing for simultaneous multi-region neural recording^1–3^. Integration of miniaturized photonic components with electrophysiology allows for localized optogenetic modulation^4–6^. Simultaneous recording of neural activity and various physiological parameters has led to new discoveries regarding the neurological effects on intracranial temperature^7^, pressure^8^, chemical concentration, etc. Combination of various functionalities, by integrating optical fibers^9–13^, μLEDs^5,14–16^, temperature sensors^17^, drug delivery channels^18^ and chemical sensors^19^, can open up a more comprehensive study of the neuronal physiology. The conventional approach for channel scaling and multifunctional integration is through monolithic integration by taking advantage of advanced micro/nanofabrication. However, the flexible probe bodies, commonly built from polymer thin films, introduced significant scaling challenges because compatible material and process options are limited. Increasing the number of polymer and metal layers may deteriorate adhesion failures. Co-fabricating other functional components monolithically on the same substrate may also result in compromised performance and yield.

Alternative to the monolithic fabrication approach, we present an innovative self-assembly approach to overcome these challenges. Our approach is to separately fabricate individual flexible probes, which serve as “building-blocks,” and stack them on top of each other into a single body in an aqueous solution by magnetic and capillary forces, which we have denominated as “hybrid self-assembly” (Figure 1a). The number of recording sites can be easily extended by stacking multiple recording probes (Figure 1c, left), and various multi-modality probes can be built by assembling other functional probes (Figure 1c, right) in a modular way. Our assembly approach enables high-precision self-alignment between the probes (less than 20-μm longitudinal, and 5-μm transverse misalignment). Also, we can achieve a high throughput, compared to manual lamination of devices using a mold^20^ or a microscopic aligner^21^. The multi-modality probes produced with this approach are dubbed as magnetic μ-laminated intracortical interface (MULTI, Figure 1a). MULTI probes with a variety of modalities and dimensions were designed with a shank width of 70∼130 μm and a thickness of 8∼15 μm. The length can be extended to meet the needs of applications and is typically 5∼10 mm. The possibility of adaptively selecting the desired functions and different configurations allows for high versatility and flexibility in various applications.

**Figure 1.**
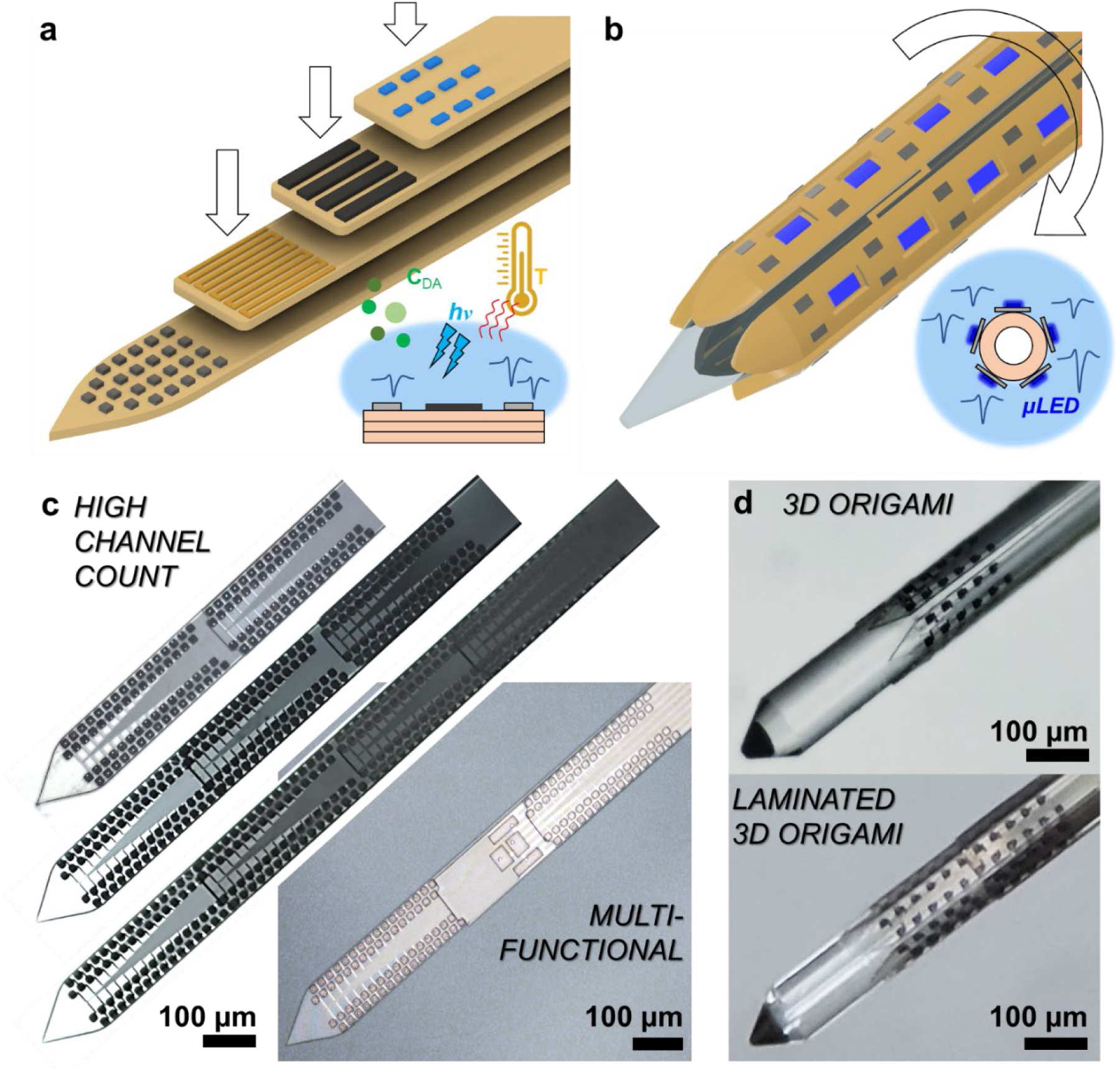
Origami flexible neural probes created by **a.** stacking and **b.** wrapping. **c.** Probe shank photos of high-channel-count and multifunctional MULTI. **d.** 3D origami probes wrapped around a sharpened optical fiber (top) and a stacked 3D origami probe (bottom). Scale bar, 100 μm.

Extending the field of view of a recording electrode or a neural probe is another important aspect of enhancing recording capability since it increases the effective recordable volume of neurons, hence resulting in a higher single unit yield^22–23^. The conventional approach for building a neural recording electrode with a wide “field of view” is by forming them at the tip of a needle-shape structure, such as Utah arrays, microneedles electrodes^24–26^, and fiber electrodes^27–28^; or by exposing multiple sides of a planar structure, such as multi-sided electrodes^29–31^, free-standing nanostripes^32^, and edge electrodes^33–34^. However, with the electrodes built at the needle-tips only, the site count and span in the depth direction is limited. As for the multi-sided electrodes approach, the fabrication and assembly are significantly more challenging. The mechanical robustness and long-term stability of such partially or entirely free-standing thin-film electrodes have not been fully established. Lastly, building three-dimensional configured electrode arrays by monolithic fabrication^23–26^ or manual assembly^27–30^ are also highly challenging and constrained when attempting to miniaturize the device size or increase site count and density.

To address the limitations and challenges of conventional approaches, self-assembly approaches have recently been explored to build three-dimensional configured electrode arrays^35–40^. For example, Neurotassel and its subsequent development^37–40^ utilized capillary force to self-assemble the microelectrodes patterned at the tips or along the length of thin-film polyimide stripes. In these previous published works, a self-assembled electrode array with a 360° field of view for 3D recording was achieved by wrapping the flexible neural probes around a circular optical fiber. The complete field of view achievable through this method provided improved recording and mapping capability compared to single-sided electrode arrays. In our implementation, we introduced ultra-compliant elastic hinges of the foldable structures by optimizing the hinge density, shape, and thickness. We were able to build a 3D origami probe with higher-density electrodes in a miniaturized dimension (a cross-section diameter of 80∼125 μm). More importantly, the ultra-compliant hinges enabled us to fold stacked probes to realize multi-modality origami probes, which effectively combined modular integration and 3D configurations. Specifically, the hinges were designed to be stretchable so that no drift or delamination between stacked probes occurred during folding. We demonstrated it by adding a “3D origami µLED module” to the “3D origami recording module” and built a 3D µLED origami probe (Figure 1b). With this 3D origami µLED module, a unique state-of-the-art spatial resolution of optical stimulation with angular selectivity was achieved by illuminating the individually addressable µLEDs distributed 360° around the core. Meanwhile, the stacked 3D origami recording module enabled in-situ electrophysiology recordings during the optogenetic modulation.

## 2. Results

We exploited the low Young’s modulus and high yield strain of polymer thin films for bending and folding into or around a desired shape in our 3D origami assembly. When emerged from liquid, the low mass-density and large surface-to-volume ratio of polymer films make them easily deformable with minimal forces. Based on these properties, we developed two self-assembly approaches. The first approach utilized magnetic forces to self-assemble multiple flexible probes into one MULTI device. The second approach utilized capillary force to wrap the device around a fiber core (Figure 1b). The same approach can also be used to fold device into a planar double-sided probe (Figure S1). For demonstration of feasibility, a variety of origami probe arrangements were fabricated, assembled, and tested, as shown in Figure 1d. Each probe has one implantable probe shank where the functional components, such as electrophysiology recording sites, temperature sensors, and dopamine sensors, as well as µLEDs, are located. This shank is extended through a monolithically fabricated flexible cable to a backend where ball bonding pads are placed for assembly to a printed circuit board (PCB). A photo of the whole probe is shown in Figure 2d. Examples of the probe shank layouts are shown in Figure S2. Polyimide (PI-2610, HD Microsystems) thin films are used as the flexible substrate and encapsulation for each probe. Interconnection metal traces were patterned with a minimum pitch of 1-μm/1-μm (linewidth/spacing) on the probe shank to ensure a miniaturized probe width while accommodating high-site-count and high-density recording electrodes (up to 64 per shank). The recording electrodes were made of surface-roughened Ti/Pt. The roughening process developed in our previous work^41^ significantly reduced the electrode impedance while enhancing electrode-to-substrate adhesion. The metal thermistor was made of 100/1000/100 nm Ti/Au/Ti in a meander pattern. Dopamine sensing electrodes differed from electrophysiology recording sites in electrode size and surface functionalization. The µLEDs were fabricated on GaN-on-Si wafers, as reported in our previous publication^16^.

**Figure 2.**
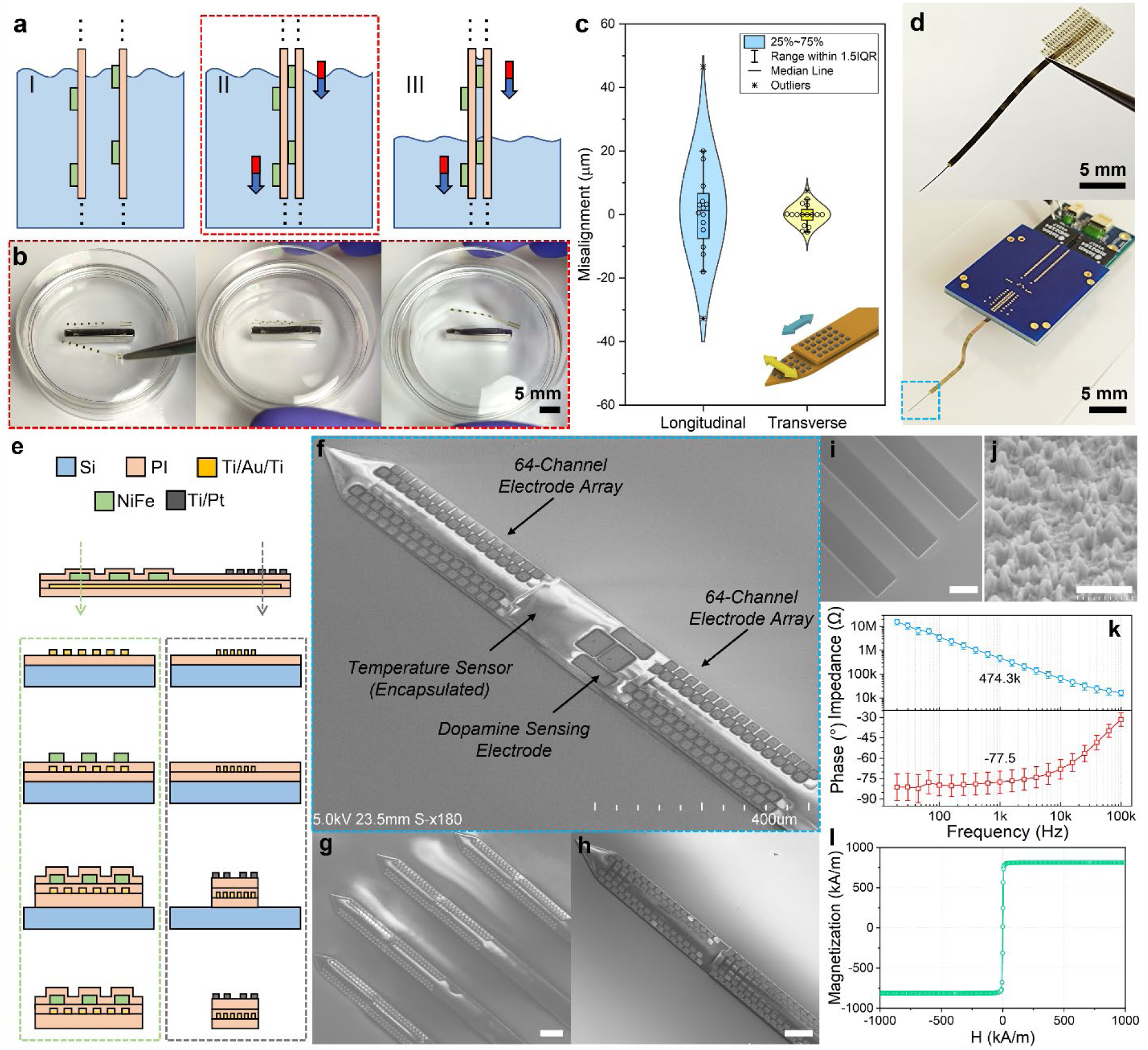
Magnetic-assisted hybrid self-assembly and MULTI probes. **a.** Illustration of the three-step assembly process: loading the probes in IPA, magnetic coarse alignment, and capillary fine alignment. **b.** Demonstration of magnetic self-alignment in which two probes with an array of magnets are attracted and aligned in IPA. The external magnetizing field was provided by a permanent magnet located at the bottom. **c.** Longitudinal and transverse misalignment tolerance in the self-assembled probes (N=15). **d.** Photograph of a MULTI probe before and after packaging on a PCB. **e.** Simplified fabrication processes for a recording probe with an array of micro-magnets plated on the cable extension. SEM photographs of **f.** a multi-functional MULTI, **g.** a 512-channel, 4-shank MULTI probes assembled by stacking two four-shank probes, **h.** a 128-channel, single-shank MULTI probe, **i.** Ni_80_Fe_20_ micro-magnets, and **j.** Roughened Pt electrode surface**. k.** Electrochemical impedance spectrum of the roughened Pt recording electrodes (N=16). **l.** Hysteresis curve of Ni_80_Fe_20_ magnets. Scale bar in **g.** to **i.** 100 μm, **j.** 1 μm.

### Magnetic-force-assisted hybrid self-assembly and MULTI

MULTI can achieve high versatility and scalability by stacking separately prepared flexible probes into one. The magnetic-force-assisted hybrid self-assembly approach enables stacking multiple probes with precise alignment at high-throughput. We used an array of Ni_80_Fe_20_ micro-magnets with a size of 100-μm by 500-μm and a thickness of 4 μm. These small magnets were electroplated only on the flexible cable, not on the probe shank. During the self-assembly process (Figure 2a), the attraction force between the arrays of magnets on the cables of the two probes could bring the two probes close to each other and then self-aligned in isopropyl alcohol (IPA) with the presence of an external magnetizing field. The external field was applied in parallel with the probes to magnetize the magnets to saturation. This step is shown in Figure 2b and the supplement movie S1. A pair of probes were placed in the same orientation in IPA. Next, the IPA container was shaken gently in the direction perpendicular to the probes with a travel distance of ∼5 mm, so that the probes could settle in the minimum energy position. Coarse alignment was achieved as the arrays of magnets were attached head-to-tail in IPA. Next, the probes were pulled out of the IPA. During this process, the probe shanks were further aligned and attached together by capillary force, achieving fine alignment.

Misalignment tolerance in the proposed self-assembly was evaluated from the 16 pairs of mock-up probe samples. These mock-up probes are identical to the actual functional probes in terms of material, design, dimensions, and magnet arrangements, but they do not have the backend electrical connections. We examined the longitudinal and transverse misalignment between the two probes after self-assembly. Except for two outliers, we obtained good misalignment tolerance of less than 20-μm in longitudinal direction and 5-μm in transverse direction, respectively (Figure 2c). Besides, this self-assembly approach could be applied for 3D origami probes (Figure 1d) and multi-shank flexible probes (Figure 2g). Once the MULTI probes were assembled and fully dried, they did not separate even when re-immersing in IPA for an hour. The adhesion between the two probes was quite strong, but not completely irreversible. If rework is necessary, friction force can be applied on its surface using a tweezer to separate the stacked probes without damage. By repeating the self-assembly process, more probes can be added to the top to integrate more modalities. For example, by stacking two 64-channel recording probes and one dual sensing probe, which is composed of one temperature sensor and four dopamine sensing electrodes, we can assemble a multifunctional MULTI device with 128 recording channels and two additional sensing modalities (Figure 2f). Each functional layer has a thickness of 5um. Therefore, the total thickness of the fully assembled MULTI device was 15 μm. Figure 2d shows the multifunctional MULTI before and after ball bonding to a PCB. The PCB was assembled with matching connectors to an Intan 128-channel amplifier board for signal acquisition. The entire backend module (with an amplifier board connected) had a weight of 5.8 grams and a size of 24 mm by 39 mm.

The simplified schematic microfabrication process is illustrated in Figure 2e (more details are given in the methods section). Figure 2f shows a SEM image of the probe tip of the MULTI probe. The metal thermistor temperature sensor was encapsulated in polyimide and therefore, not visible under the SEM. Four interconnection lines were connected to the thermal resistor to accurately measure the resistance change using a four-point measurement. The temperature coefficient of resistance was 0.19 ± 0.014% (N=4, Figure S3). The roughened recording electrodes were formed by sputtering Ti/Pt over the roughened polyimide surface. The surface morphology was visualized in a SEM image at a 45° angle (Figure 2j). At 1 kHz, the electrochemical impedance was measured as 474.3 ± 34 kΩ (N=16 electrodes) from an LCR-meter (Keysight E4980A) in phosphate-buffered saline (PBS 1M). The roughening process reduced the electrode impedance by three-fold. The DC magnetic hysteresis curve of permalloy micromagnets was measured using a vibrating sample magnetometer (VSM) (VersaLab™, Quantum Design). The resulting M-H curve was diamagnetically corrected and shown in Figure 2i, indicating a magnetic saturation at ∼813 kA/m.

### High channel count scalability by self-assembly

The ability to scale channel count can be achieved by simply stacking multiple recoding probes. Figure 1d illustrates that four single-shank flexible probes (each with 64 recording electrodes spanning 400 μm) could be stacked by self-assembly to create a 256-channel probe, which is to the best of our knowledge, the highest channel count for a single shank flexible probe. The width and spacing of interconnection traces were both 1μm. The area of each recording electrode was 250 μm^2^. The recording site’s center-to-center pitch was 25-μm between two adjacent electrodes for high spatial resolution. Even a higher channel count of 512 could be achieved by stacking multi-shank probes, as shown in Figure 2g. For a proof-of-concept demonstration, we only recorded signals from 128 channels. The total number of simultaneous recorded channels could be further extended by integrating application-specific integrated circuit (ASIC) chips, as demonstrated in our previous work^41^.

### Simultaneous neural recording and local tissue temperature monitoring

We used a multifunctional MULTI probe (Figure 2f) that includes 128-channel neural recording, temperature monitoring, and dopamine sensing. Urethane anesthesia was used in this acute experiment in a rat. The MULTI probe was implanted to target simultaneous recording in cortex and hippocampus regions. The probe shank was mounted on the surface of a 200-μm diameter stainless-steel syringe needle and coated with PEG. The syringe was fixed on a linear manipulator for insertion. Details of the anesthesia, surgery, and probe insertion are described in the method section. The syringe needle was kept un-retrieved after delivering the probe to the target brain region.

We first performed a 30-min neural recording simultaneously in both cortex and hippocampus. Wide-band signals (0.1∼7500 Hz) with a duration of 100 ms are shown in Figure 3a. The waveforms, autocorrelation histograms, amplitude, and trilateration locations of the single units are shown in Figure 3b, and c. Thirty different single units were observed and distinguished along the two 64-channel electrode arrays located above and below the temperature and dopamine sensors in the middle. Next, the local brain temperature was modulated by heating and cooling the stainless-steel needle implanted with the probe. The resistance of the temperature sensor was continuously measured using a source-meter (Keithley 2400) in a four-point configuration. The applied DC current during sensing was 10 μA. The resistance of the sensor at 37℃ was characterized as 15.76 kΩ in bench top tests. The sensor self-heating induced by the driving current was 1.576 μW, which would not cause any detectable temperature increase (<0.01 ℃) during the 15-min baseline measurement. Next, tissue temperature was modulated from the control period (no-modulation) to cooling and heating periods. Temperature was recorded every 2 minutes. The change of neuron firing rates were recorded as the brain temperature changed over time (Figure 3d). For each neuron, the z-scores were calculated from the average and standard deviation from the first 15-min recording (control period). As shown in the heatmap, the firing rate of 19 pyramidal neurons (out of 24) showed positive correlation with temperature. The average firing rates for the pyramidal and interneuron populations are shown with red lines on top of the heatmap (Figure 3d). In contrast, some interneurons showed significant firing rate reduction during the heating period as well. During the presented temperature modulation period, a positive correlation between sharp-wave-ripple occurrence rates and temperature was observed. The average ripple frequency during the cooling period decreased (129.1 Hz) from the one during the control (133.4 Hz) and heating (133.2 Hz) periods. Same trends of firing rates and average ripple frequency changes were observed in the other two repeated trials. Similar phenomena were also reported in previous research^42^.

**Figure 3.**
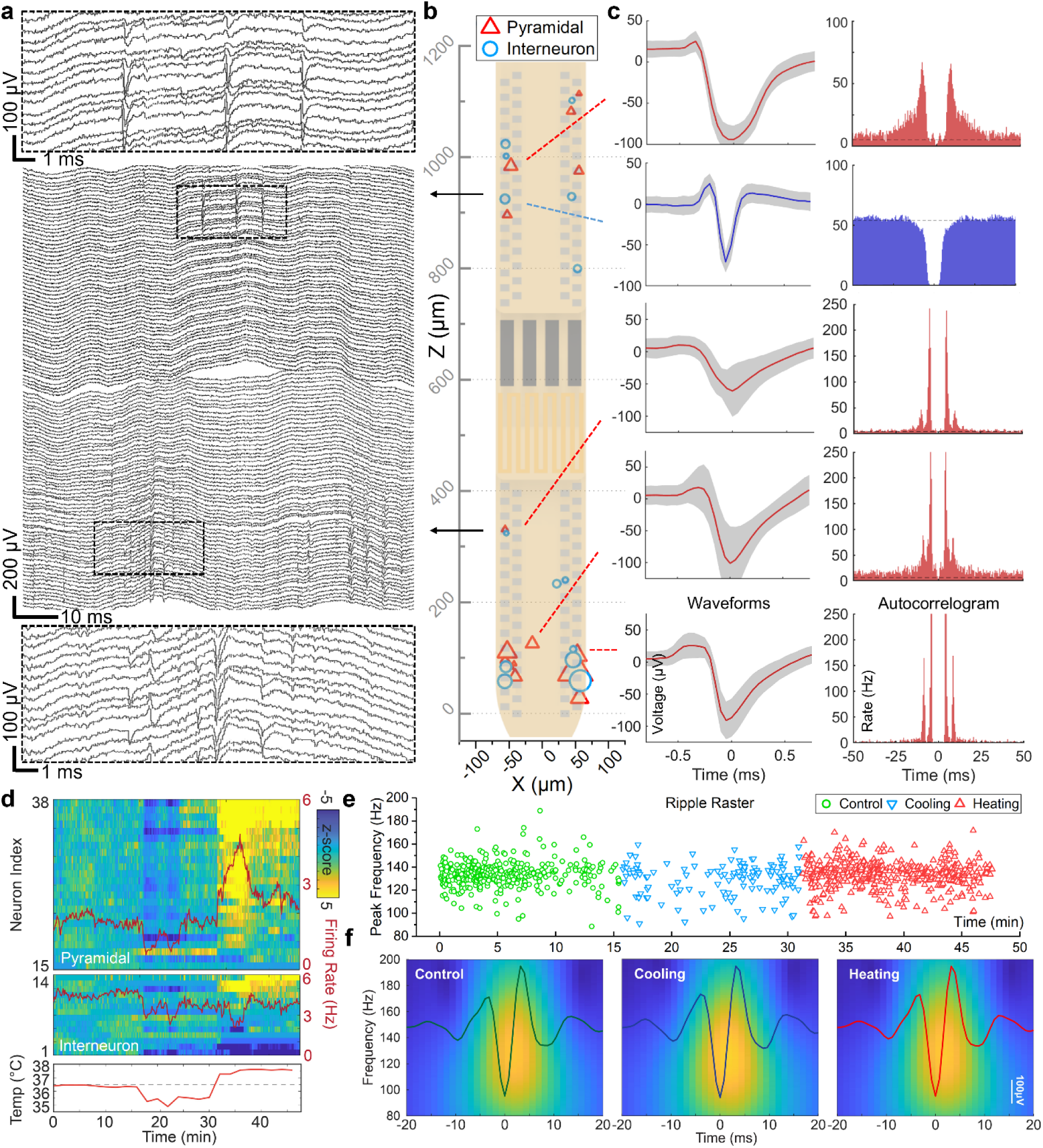
Simultaneous neural recording and local tissue temperature monitoring using a multifunctional MULTI probe. **a.** Wide-band signals recorded from 128 recording sites. Dashed rectangles show a zoomed-in view to illustrate spiking activities in both cortex and hippocampus. **b.** Probe layout shown with the putative locations of recorded neurons. The symbol size represents the spike amplitude with an average value of 71.5 μV. **c.** Putative pyramidal cells (red) and interneurons (blue) were recorded from both regions. Representative waveforms and auto-correlation histograms are shown on the left and right, respectively. **d.** The average (red) and normalized (z-score heatmap) firing rates of pyramidal neurons and interneurons aligned with the temperature change. **e.** Local temperature manipulation affected the peak frequency and the occurrence of ripples. Cooling reduced the number of ripples (blue triangles), while heating increased it (red triangles). **f.** Mean ripple waveforms and wavelet maps of ripples detected during the three different temperature conditions. Heatmap scale, 0 to 1500 dB.

### Simultaneous neural recording and dopamine concentration monitoring

Dopamine sensing was achieved by amperometry: measuring currents generated by oxidation of dopamine from an electrode at oxidation potential of 0.4 V (with respect to the reference electrode of Ag/AgCl 3M NaCl). To increase the dopamine detection sensitivity, we modified the Pt electrode surface with reduced-graphene oxide (rGO). The electrochemical properties of the reduced-graphene oxide (rGO) electrodes were tested with electrochemical impedance spectrum (EIS), cyclic voltammetry (CV), and amperometry. Details of the measurement setups and procedures are described in the Methods section. Nyquist plots obtained by EIS measurement before and after rGO modification showed that the electron transfer resistance was reduced from 385.9 kΩ to 143.0 kΩ. With rGO modification, the peak current of cyclic voltammogram increased from 114.1 nA to 334 nA (Figure 4a). The dopamine detection sensitivity and selectivity were characterized with benchtop tests by introducing dopamine and other interference chemicals with known concentrations to PBS during amperometry measurement (Figure 4b and d). The sensitivity was measured as 286 ± 12 pA/μM with a linear range from 2.4 μM to 153.4 μM and a detection limit of 28 nM (Figure 4c). The selectivity test showed that γ-aminobutyric acid (GABA) and uric acid (UA) gave negligible effects while amino acid (AA) gave a slight (<10%) current increase (Figure 4d).

**Figure 4.**
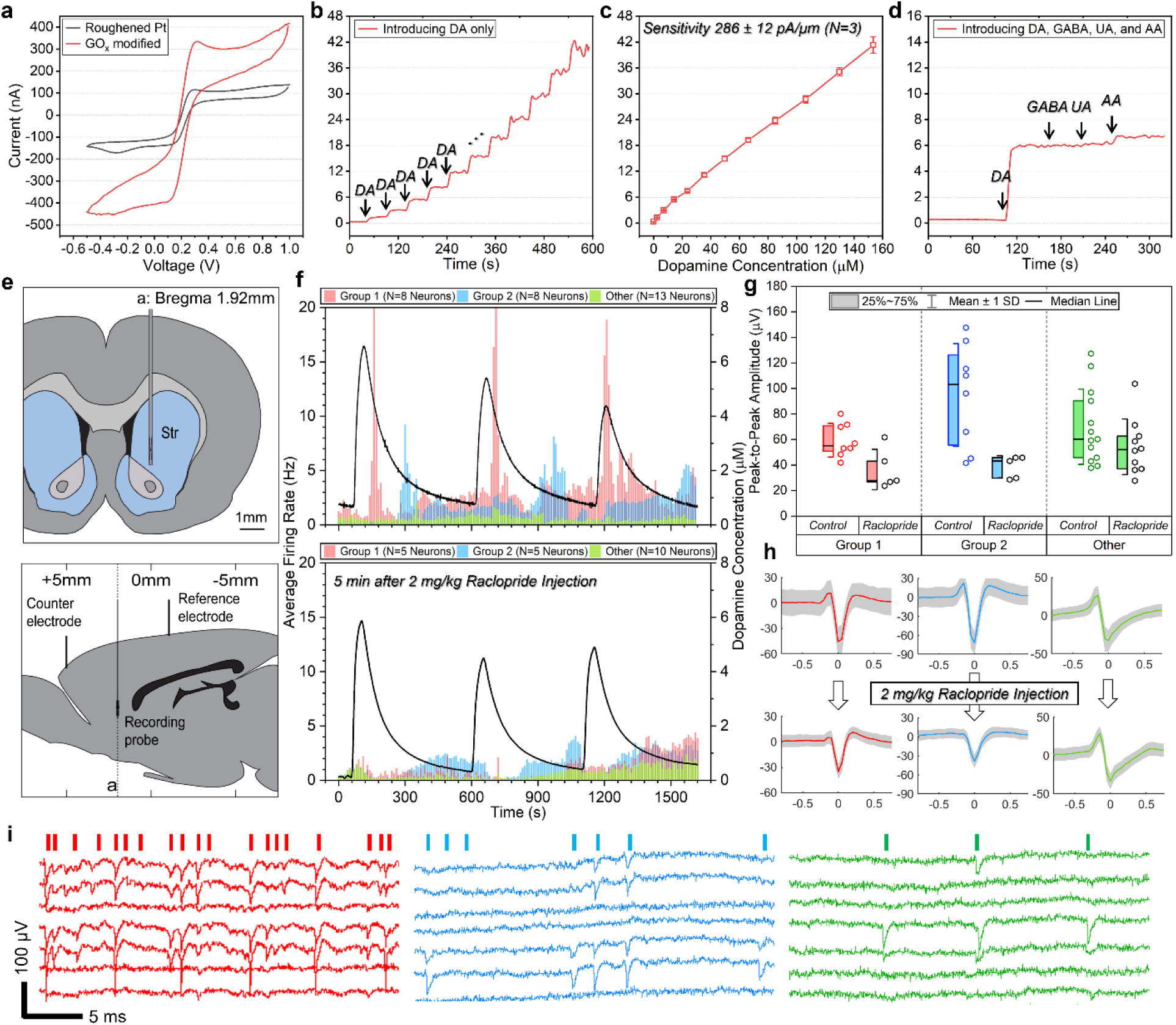
Simultaneous neural recording and dopamine concentration monitoring with the multifunctional MULTI device. **a.** Cyclic voltammetry of the electrode before and after rGO surface modification. Amperometry measurement of the dopamine sensor during **b.** sensitivity and **c.** The calibrated sensitivity of the dopamine sensor using amperometry (N=3). **d.** selectivity tests. **e.** Implantation position diagrams of MULTI (mounted on a stainless-steel needle), Ag/AgCl counter and reference electrodes. **f.** Average firing rates correlated with the local dopamine concentration of the three groups of neurons clustered based on their firing patterns before 2 mg/kg Raclopride injection. **g.** Mean amplitudes of all the single units and **h.** mean waveforms of the representative single units before and after Raclopride injection. **i.** Wide-band signals with a duration of 30 ms zoomed-in to the spikes of the neurons from each group.

Simultaneous recording of neural activities and local dopamine concentration measurement was performed from the dorsal striatum in an anesthetized rat. The MULTI probe was mounted on a stainless-steel injection needle (190 μm in outer diameter) for insertion. The needle was left in place for dopamine injection into the striatum. Details of the implantation and dopamine measurement procedures are described in the Methods section. We injected 400 nM dopamine solution to induce a peak local dopamine concentration between 1 to 10 μM at the middle of the probe where the dopamine sensor was located. From the recording electrodes, 29 different single units were identified before an intraperitoneal injection of Raclopride (2 mg/kg) while local dopamine concentration was monitored simultaneously. Benefiting from the high-channel-count recording capability of MULTI, various types of neuronal behaviors could be identified and studied. Before Raclopride injection (control), the neurons were clustered into three groups by their firing patterns (Figure 4f). More details are described in the Methods section. We also observed that the clustering was uncorrelated to the trilaterated soma locations. Starting 5-minute post-Raclopride injection, the same neurons of each group were identified based on the soma locations and autocorrelation histograms. Significant reduction of the peak amplitude and firing rate was observed among group-1 and group-2 neurons (Figure 4f, g and h). A few neurons of each group were no longer detected due to either a complete suppression of the activities or a reduction of the amplitude to below the noise level (∼20 μV peak-to-peak).

### Capillary assisted self-assembly of stacked 3D origami probes

The controllable capillary assisted wrapping of the origami neural probes was enabled by ultra-compliant hinge designs (Figure 5a, b). The hinges were defined by a two-step reactive ion etching as detailed in Figure 5a. The first etching step is to define the hinge shapes by completely removing most of the polyimide and leaving the elastic hinges connecting the sub-shanks. The second etching is for further thinning the hinges to make them ultra-compliant. The hinges provide mechanical connection to sub-shanks holding the shape but are compliant to be bent in a controllable manner during the self-assembly. Various 3D origami probes with different shapes were fabricated by adjusting the design and total width of the shank to fit to a specified fiber diameter. As a demonstration, three types of 3D origami probes were made to wrap around the fiber cores with different diameters of 125-μm, 105-μm, and 80-μm, respectively.

**Figure 5.**
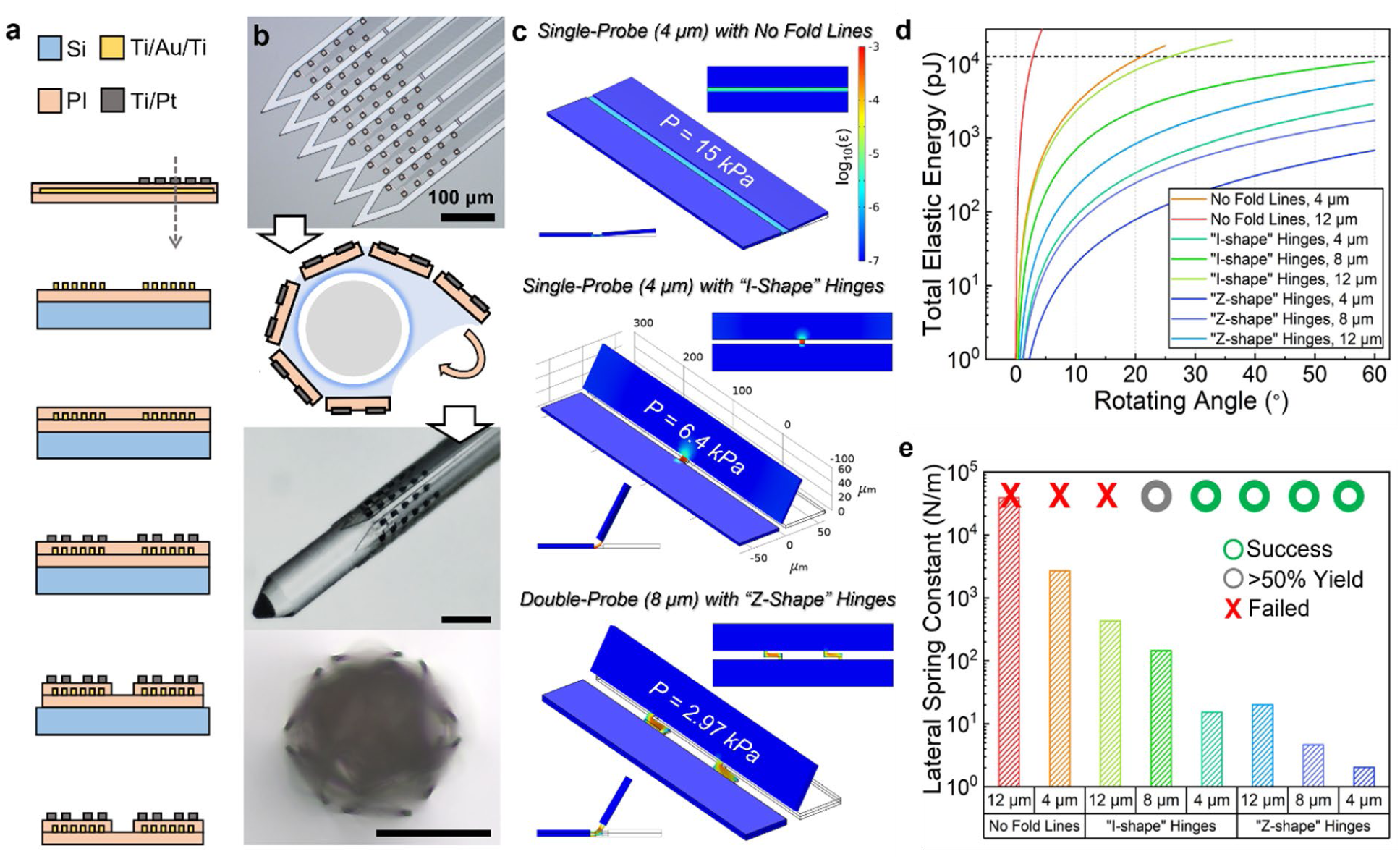
**a.** A simplified fabrication process summary for 3D origami probes. **b.** The tip of an origami probe before being released from the wafer (top), and the side and cross-section view during and after wrapping (bottom). Scale bar, 100 μm. **c.** Deformation and strain distribution of several models with various elastic hinges. Hinges can be bent by over 60° when the modeled inward pressure is less than the estimated capillary induced pressure. **d.** The total elastic energy of the hinges. The dash-line marks the estimated surface energy reduction. **e.** The lateral spring constant at small deflection and experimental results.

We first immersed the fabricated probes in IPA, which has a lower surface tension coefficient compared to water. With the specially designed elastic hinges, the probes remained flat during and after being removed from the IPA. After being fully dried, the probe was transferred to a temporary holder with its shank free standing. Next, an optical fiber was coarsely aligned and attached at the end of the flexible cable with its tip protruding. Finally, the free-standing probe shank and the fiber tip were immersed into DI water and then slowly (∼1 mm/s) pulled out. During the pulling, the origami probe sub-shanks were fully wrapped over the fiber by capillary force. A video of this process is provided in the supplementary (movie S2). In addition to a single layered 3D origami probe, we could assemble multi-layered 3D origami probes for channel count scaling and multifunction integration. To achieve this, the two planar origami probes with embedded micro-magnets were first self-assembled in IPA, then wrapped around a fiber, following the same pulling process in DI water.

To obtain successful wrapping, we simulated and experimentally validated various types of hinge designs with different thicknesses. During the wrapping process, the elastic energy of hinges increases as the structure deforms, while the surface energy decreases as the probe is pulled out of DI water. The inward pressure on the sub-shanks induced by capillary force increases as the liquid exits and the gap between the probe and fiber diminishes. Based on the geometry of the shanks (sub-shank width of 60 μm, hinge pitch of 400 μm), the total surface energy reduction and maximum inward pressure achievable were estimated to be 13.9 nJ and 15 kPa, respectively. Details of the analytical method for estimating the elastic energy and capillary forces are presented in the supplementary (Figure S4). These results were used as a threshold for predicting successful wrapping with 60⁰ hinge bending. Following this analysis, control (“no fold lines) and two separate types of hinges (“I-shape,” and “Z-shape”) with various hinge thicknesses were analyzed to evaluate feasibility of the self-assembly process for 3D origami probes. The results are shown in Figure 5c-e. Based on the maximum inward pressure achievable with capillary forces, one failed (top) and two successful cases (middle and bottom) are shown in Figure 5c. The total elastic energy of the hinges at different bending angles were simulated and plotted. The dash-line indicates the total surface energy reduction estimated using the analytic method, which corresponds to the threshold for successful wrapping. The lateral spring constants for different hinge types and thicknesses are modeled and the experimental wrapping results are indicated in Figure 5e. The net system energy change combined with the modeled inward pressure required for 60° hinge bending could be a good indicator of whether the wrapping could be reliably achieved for various shank and hinge designs. After wrapping, the origami probe remained in good attachment on the surface of a fiber in a hexagonal shape. We also verified that the 3D origami probes did not detach from the fiber during insertion. The tests were performed by inserting the assembled probes (N=3) into a phantom brain filled with agarose gel (0.6 wt.%), which mimics brain tissue. The tips of the probes were imaged perpendicularly to the insertion direction using a transmission microscope. The schematic of the setup and the cross-section images taken at different insertion depths are shown in the supplementary (Figure S5). Even without any adhesive materials applied, no delamination or unwrapping was observed during the insertion and retrieval tests. Each time, the probe was inserted into the phantom brain over 8 mm deep.

### High-density three-dimensional neural recording

For acute recording, we prepared a 3D origami probe of six sub-shanks containing total 64 recoding sites (Figure 5b). First, a fiber (105 μm in diameter with a tapered tip) was first mounted onto a PCB holder, which was used to fix the PCB onto the linear manipulator for insertion. The position of the flexible cable was adjusted to align the origami probe shank with the fiber. The probe and the optical fiber were then self-assembled in DI water and ready for insertion. To further secure the probe onto the fiber, PEG coating can be applied. The cone-shaped tip of the fiber was sharp enough so that no additional insertion shuttle was needed.

Single unit recording from neurons 360° surrounding the 3D origami probe was successfully accomplished. The mean spike amplitude and soma locations are shown in Figure 6a. Trilateration of the neurons from 3D recording was performed using a 12-column electrode array model (2 columns on each sub-shank) with periodic boundary condition. The algorithm was based on the open-source package CellExplorer^43^ with minor adjustments described in the Methods section. The trilateration planar coordinates obtained from CellExplorer were converted into polar coordinates to calculate the approximate angular positions and z-coordinates of the neurons but the perpendicular distance from the source to the 3D probe surface was not determined (Figure 6a and b). The probe illustrations in the figure are slightly reduced in size for better visualization. Angular heterogeneity can be easily observed in both single unit activities and local field potentials, as shown in the waveforms, spike raster plots, and the representative single units. Expanding the field of view to 360° contributed to holistic mapping of neural circuit connectivity. In comparison, the conventional single-sided probes have huge blind spots, almost a half of hemispheres on the backside of the probe where no recording sites are present.

**Figure 6.**
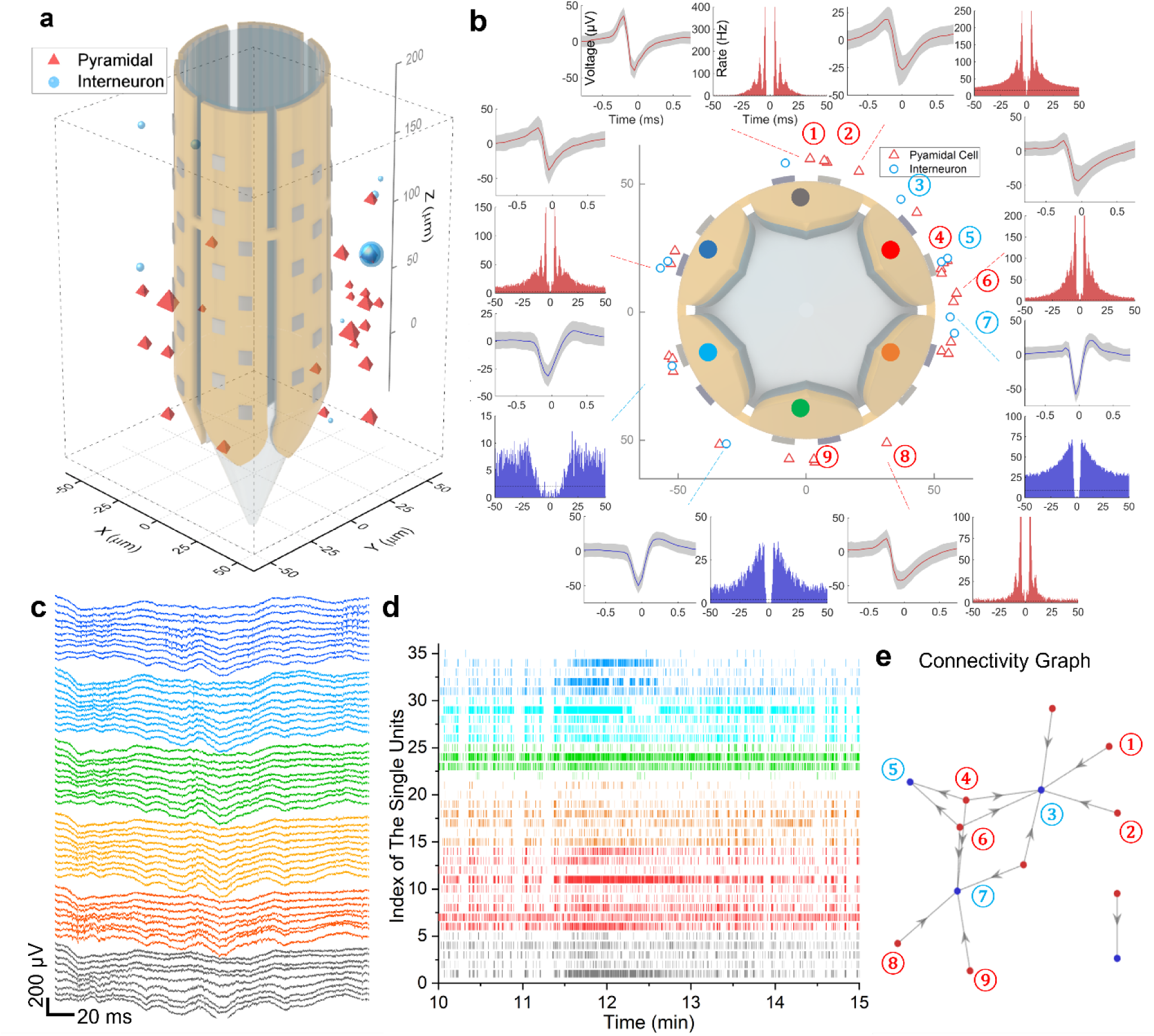
Acute recording with a 3D origami probe. Single units identified surrounding the 3D probe and their projected positions on the probe surface in **a.** 3D and **b.** top view. The symbol size in **a.** represents the spike amplitude with an average value of 95.3 μV. The probe illustrations are not to the scale. **b.** Autocorrelation histograms and mean waveforms of representative units. **c.** Wide band (0.1-7500 Hz) neural signals recorded using 3D origami probe. Signals from each sub-shank is annotated with the same color. The corresponding locations are marked in **b.** Same color coding is applied to **d.**, the spike raster plot during a representative 5-min recording period. **e.** Neural circuit identified surrounding the probe after detecting putative monosynaptic connections between neuron pairs (details in Methods section).

### Acute three-dimensional opto-stimulation and electrophysiological recording

To achieve directional optogenetic stimulation and electrophysiological recordings, a multi-layered 3D origami probe was prepared. A 3D origami recording module of 5 sub-shanks with 10 Pt recording sites on each shank was stacked on top of a 3D origami μLED module with 4 blue GaN-μLEDs per sub-shank. Together, the multi-layered stacked optoelectrode was wrapped around a fiber of 80 μm in diameter using the assembly method previously described (Figure 7a). The assembled 3D µLED origami probe was then covered by PEG and inserted into the dorsal hippocampus of an anesthetized Thy-1 transgenic mouse as shown in Figure 7b. Each µLED was individually addressed and sequentially triggered 50 times at 2Hz, 50ms pulse duration and 2 different intensities (0.1 and 1.57 µW optical radiant flux). Spike sorting and cell body approximation was calculated, as described in the 3D recording experiment.

**Figure 7.**
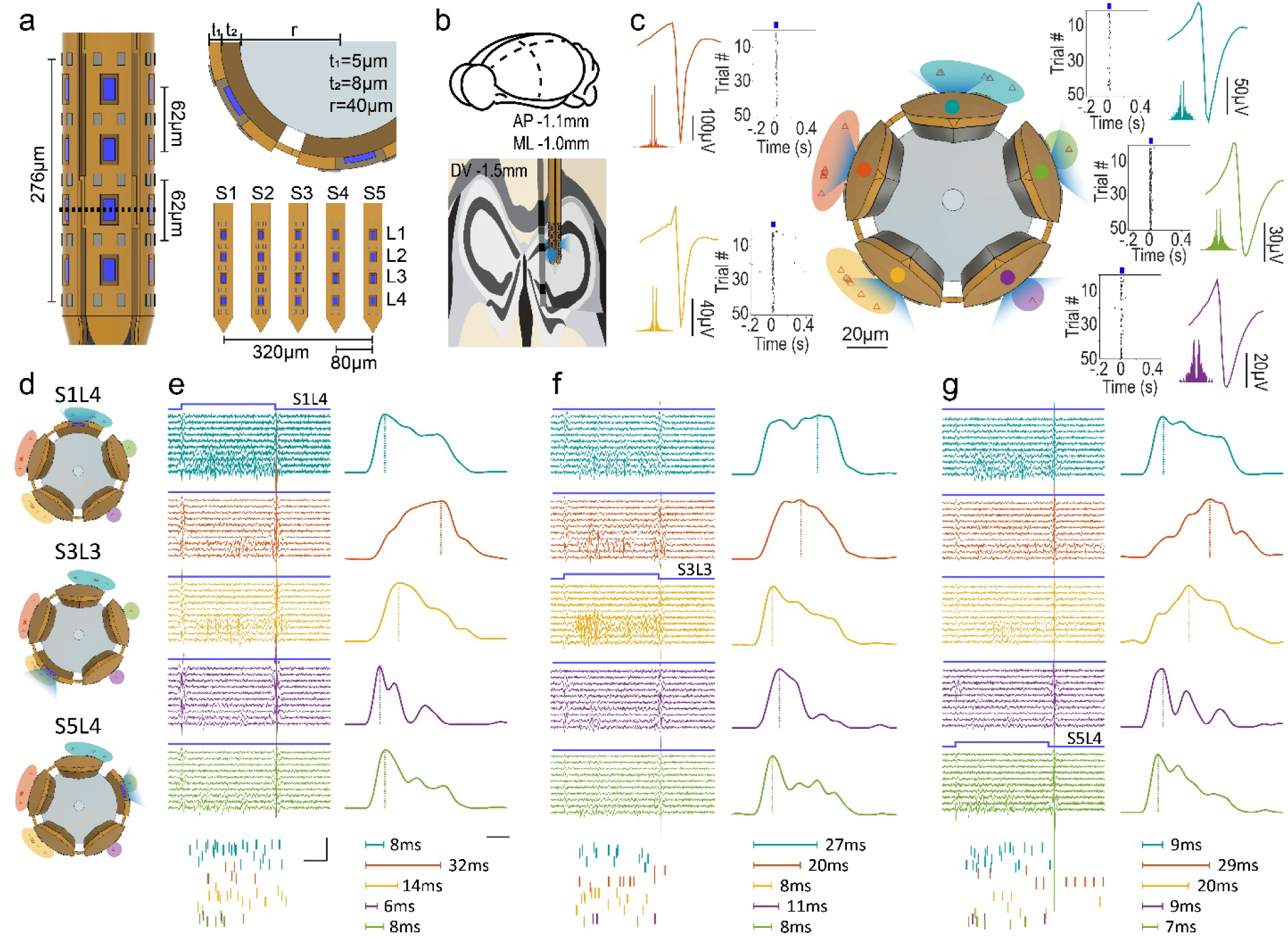
Acute directional extracellular recording and opto-stimulation with a 3D µLED origami probe. **a.** A 5 sub-shank 3D µLED origami probe with a recording-stimulation coverage of 276µm vertically per shank was wrapped around a fiber of 80um in diameter. **b.** Location of the implanted probe in the dorsal hippocampus of a Thy-1 transgenic mice (AP-1.1mm, ML-1.0mm and DV-1.5mm). **c.** Directional optogenetic stimulation resulted in an increase in the firing rate in neurons facing each sub-shank. Waveforms and raster plots are color coded to match their corresponding shanks. **d.** Representation of the cross section of probe, the illumination cone is shown for each of the results shown in e-g. **e-g.** Directional optogenetic stimulation evoked local and non-local increase in the population’s firing rate with different latencies depending which μLED was triggered (**e.** S1L4, **f.** S3L3, and **g.** S5L4). Bandpass filtered (300-6000 Hz) traces from each shank (scale: 10 ms by 25 µV), raster plots and PSTHs (scale: 10 ms), and color coded to match the units enclosed in the shaded area facing each shank. The vertical dashed line marks the population’s maximum.

A total of 28 single units were isolated from the recording near the tip of the probe (26 putative pyramidal neurons and 2 putative interneurons): 8 on shank 1, 7 on shank 2, 10 on shank 3, 1 on shank 4, and 2 on shank 5. Directional lighting increased the firing rate of 27 neurons out of the total 28 single units in response to at least one of the triggered µLEDs (Figure 7c). For most cases, more than one µLED induced a significant increase in the firing rate of a particular unit. Figure 7d-g shows the shank’s neural dynamics in response to three different µLEDs (S1L4, S3L3 and S5L4). High-pass filtered traces (>300Hz) are shown for a single activation of S1L4, S3L3 and S5L4 (Figure 7e-g). Clear neuronal responses can be observed in multiple shanks for all 3 µLEDs. The raster plots below the traces confirm the evoked action potentials during the light period (50ms pulse). Peristimulus time histograms were plotted for each shank by binning the APs from the corresponding neurons in each shank (in 1ms bins, APs within -2 to +3 ms to the LED onset were discarded), aligned to the µLED onset, normalized to the maximum value in each shank and smoothened with a 10ms gaussian (full list of PSTHs for both power outputs are shown in Figure S6). Neurons in front of the triggered µLED responded within 6-8ms. Additionally, off-site and radially opposite neurons also responded within the same time frame, i.e., neurons in shank 4 responding to shank 1 illumination, or neurons in shank 5 responding to shank 3 illumination, potentially suggesting antidromic activation of the offsite neurons. In contrast, longer latencies were observed for the neurons in shank 2 and 3 when those in neighboring sub-shanks or radially opposite shanks were triggered, likely due to polysynaptic activation. These results imply the potential of the 3D µLED origami probes for studying local circuitry of an intricate structure like the dorsal hippocampus.

### Double-sided neural probes

The double-sided probes can be made by reducing the number of sub-shanks to two from the 3D origami recording module (Figure S1a). The number of hinges should be minimum to make it super compliant to be folded into a double-sided probe. The assembly process is similar to the 3D origami probes but without fibers. A double-sided probe was tested with short-term implantation to the rat’s dorsal hippocampus (1 week). The average peak-to-peak amplitude of signal units 1-week post implantation was found to be 26.2% larger than the average amplitude observed in the acute 3D origami probe recording experiments, which indicated recovery of the neurons. Figure S1a shows the representative single unit waveforms and auto-correlograms. Distinctive single units from the front and the back side at the same depth were identified. Single units located within 5 μm distance in z-direction near the probe tip showed different waveforms. Sharp-wave-ripples recorded from the front and back side showed a difference in both frequency and amplitude. Figure S1c shows the wideband signals recorded near the probe tip from both sides. The difference in spike signals and local field potential could be easily noticed.

### Longevity of 64-channel flexible probes for chronic recording

Flexible recording probes with two different metal interconnection pitches, 1-μm/3-μm and 1-μm/1-μm, were chronically implanted for longevity evaluation. Each of the implanted probes were made by manual stacking of two flexible probes with the same pitch and 32 recording sites each. Both electrode impedance and recording performance were monitored over time. Spike sorting and curation were performed with the same method described in the previous sections. The total unit counts, spike amplitude distributions, average firing rates, and signal-to-noise ratios were summarized after completion of spike-sorting all the recorded data. Thus, the observer bias was minimized. The electrode impedance and the recording data summary are presented in the supplementary (Figure S7).

During the chronic recording, we observed an initial failure from a decline of the impedance values (>30 %) among a small number of electrodes adjacent to each other (2 to 4 sites out of 64 sites). Such impedance decline could indicate a short circuit between those recording channels caused by liquid migration into the flexible substrates. We mark the date of such impedance decline as the starting point of a partial device failure. For the probes fabricated with 1-μm/3-μm pitch (N=2) and 1-μm/1-μm pitch (N=3) interconnections, all the recording sites were fully functional without any partial failures for at least 12 weeks and 8 weeks after implantation, respectively. The longest longevity of those probes being fully functional was 19 weeks. After the initial observation of a partial failure, most of the recording channels remained functional for approximately 4 additional weeks, until a decline (>50%) in the number of trackable single units marked the end of the probe longevity. Thus, a good longevity of 15 weeks on average was observed (Figure S7). The probes used to create MULTI, 3D origami and double-side folded probes were fabricated and packaged with the same design rules and process parameters. Therefore, the longevity of all the probes presented in this paper is expected to be similar.

## 3. Discussion

The hybrid self-assembly technologies presented in this paper enabled several advanced neural probes with cutting-the-edge benchmarks in scaling electrode density, adding multi-modality, and configuring 3D electrode arrays. These approaches significantly reduce the burden of unconventional 3D microfabrication and enhance the yield and versatility of multilayer integration. Using the self-assembled stacking technology, we demonstrated immense scalability and customizability for various types of planar, 3D origami and double-sided probes.

A number of recording electrodes can be scaled by stacking multiple recording modules on a flexible substrate. For example, using a 64-ch recording probe as a unit module, we demonstrated that an electrode array could be extended to 128, 192, and 256 channels on a single shank (Fig. 1c). If a four-shank module is used (Fig. 2g), the electrode array should be easily scaled up to 1,024 channels. Depending on the target region of the brain and applications, the recording site density and span can be adaptively scaled and adjusted in the final assembly step. Therefore, various configurations can be customized for various target brain regions. We also demonstrated a self-assembled high-site-count multi-modality probe by stacking two 64-channel neural recording modules and a temperature and dopamine sensing module together (Fig. 2f). The customized span of the electrode arrays enabled simultaneous recording of a large population of single units in different regions. The location of sensing temperature and dopamine concentrations can be customized, and even multiplied by stacking additional sensing modules. Similarly, other modalities can be added such as optogenetic modulation and drug delivery. For self-aligned stacking, we integrated micro-magnets with a length of 500 μm in each module and demonstrated reliable assembly of four modules. The size and shape of micro-magnets can be further optimized (shorter magnets and wider spacing) to accommodate more modules assembled on the same substrate. The external magnetic field can be engineered for optimal intensity and field trajectory to enhance alignment accuracy.

With a combination of self-assembled multi-layer stacking and wrapping, a multi-modality 3D origami probe was constructed for electrophysiological recording and µLED optogenetic neuromodulation. This 3D origami probe is a perfect tool for brain mapping since it possesses a high-spatial-resolution and directional selective 360° interface with surrounding neurons. It has thus by far the highest density of recording sites and optical stimulation sites in a cylindrical 3D configuration. Using the 3D origami probes, 3D optogenetic modulation and mapping of neural circuits was feasible (Figure 6a, e). Directional selective optical stimulation was achieved by illuminating individually addressed µLEDs on the sub-shanks facing different angles. Longer latencies were observed for the neurons in the sub-shanks that were located radially opposite to the triggered µLEDs. The recording channels and optical stimulation sites can be further scaled by stacking more recording/µLED modules, as shown in Fig. 1d. On the backend headstage, the extended cable can be directly assembled with an ASIC chip to enable large-scale front-end computing, digitizing, and control in a more compact packaging suitable for chronic experiments of freely behaving animals^44^. Furthermore, a rigid fiber can be replaced by a softer material such as biocompatible polymer fibers or hollow cylindrical sheaths. More variation of cross-sections can be deployed by patterning the fiber with wither concave or convex shapes and increasing the number of the sub-shanks. For applications where a rotational symmetric full coverage of a 360° field of view is not critical, double-sided probes can be built by simply folding a two sub-shank version of the origami probes (Fig. S1). In comparison with the 3D origami probe, the spike signals sourced from the horizontal direction in plane with the double-sided probe may be difficult to record and locate. However, the miniaturized cross-sectional size (70∼130 μm in width and 8 μm in thickness) makes it more suitable for chronic implantation and small animal applications.

## 4. Methods

### Flexible neural probe fabrication and assembly

Flexible neural probes were microfabricated using MEMS technologies following the thin-film polyimide device fabrication processes. First, a Cr/Au/Cr sacrificial layer of 100/500/500 Å in thickness was evaporated (Enerjet Evaporator) on a 4-inch silicon wafer coated with a layer of 500-nm silicon oxide. Polyimide (PI 2610, HD Microsystems) was span over the sacrificial layer at 3000 rpm. A soft baking at 115 ℃ for 10 minutes was applied to remove the solvent. The polymerization process was completed in a vacuum oven (YES PB8-2B-CP Oven) at 350 ℃ for 1 hour. The average thickness of a single polyimide after polymerization was 2 μm. Second, a layer of lift-off resist (PMGI SF 5S) was span at 2000 rpm and baked at 200 ℃ for 5 minutes. On top of it, a layer of photoresist (SPR 220 3.0) was span at 2000 rpm and baked at 115 ℃ for 90 seconds. The 2 μm pitch patterns were created using a projective lithography tool (GCA AS200 AutoStep). After plasma descum for 60 seconds (Plasmatherm 790), Ti/Au/Ti with a thickness of 100/1000/100 Å was evaporated and lift-off patterned by fully dissolving the photoresist and lift-off resist in Remover PG (Kayaku Advanced Materials Inc.). After dehydration baking, the wafer was plasma activated with oxygen plasma (Tergeo Plus). The polyimide coating and curing process was repeated. Next, a seed layer and a Ni_80_Fe_20_ magnet layer were patterned. The details of the electroplating and magnet characterization process are described in the next section. An additional polyimide layer was coated with the same surface activation and polymerization process.

Third, an Al_2_O_3_ of 200 Å in thickness was deposited at 150 ℃ by atomic layer deposition (Oxford OpAL ALD) as a polyimide etching hard mask. A 3 μm SPR 220 3.0 photoresist was span coated and patterned to define the outline and “fold-lines” of the flexible probe. Diluted HF acid was used to etch Al_2_O_3_ to expose the polyimide underneath. The polyimide was then etched away by reactive ion etching (Plasmatherm 790). With a similar process, contact vias (3 μm by 3 μm openings) on the encapsulation polyimide layer were etched. For the origami probes, the connecting elastic hinges were further thinned down to 2 μm by timed etching. After polyimide etching, the Al_2_O_3_ hard mask was removed. A layer of sub-nanometer Ti was sputtered using a metal sputter deposition tool (Lab-18 Sputtering System) to form a non-continuous etching hard mask. The surface of the top polyimide layer was roughened by reactive ion etching through the unmasked Ti layer. Next, a Ti/Pt layer with a thickness of 100/1000 Å was sputtered and lift-off patterned with a similar lithography process to form the recording electrodes. Last, the sacrificial layer was removed in Cr etchant to release the probes from the wafer carriers.

### Ni_80_Fe_20_ micromagnet electroplating

A Ti/Au layer with a thickness of 100/500 Å was evaporated and lift-off patterned with a similar lithography process to form a seed layer. Next, a thicker (5 μm) layer of photoresist (SPR 220 3.0) was spun and patterned. After plasma descum, Ni_80_Fe_20_ micro-magnets with a thickness of 4 μm were electroplated under DC current with a current density of 15 mA/cm^2^ for 15 mins. A meshed Ni plate (10 cm by 10 cm) was used as an electroplating anode. The wafer and the Ni anode were immersed in an electroplating bath and approximately separated by 5 cm in a 2000-mL size glass beaker. Continuous stirring was applied with a magnetic stirrer. Details of chemical formula and concentration are described in the supplementary (Table S1).

### Dopamine sensing electrode surface modification and characterization

The roughened Pt electrode surface was pre-conditioned by cyclic potential scanning (-1.0 V to +1.0 V at a scan rate of 100 mV/s) in PBS using a potentiostat (Interface 1010E) with a conventional three-electrode configuration. Specifically, the working electrodes were Pt electrodes (40 μm x 50 μm) with roughened surface. The counter electrode was a platinum wire electrode. All the electrical potentials described in this section were relative to an Ag/AgCl (in 3 M NaCl) reference electrode. After conditioning, the electrodes were rinsed with DI wafer. Graphene oxide was dissolved in the DI water using an ultrasonic bath for 2 hours. Next, the electrodes were modified with reduced rGO by cyclic potential scanning (10 cycles, -1.0 V to +1.0 V, a scan rate of 50mV/s) in a mixture of rGO (3 mg / mL) and Na_2_SO_4_ (0.1 M) solution. The rGO modified electrodes were air-dried for 30 minutes at room temperature.

Amperometry and cyclic voltammetry measurement were performed using the same potentiostat. All the *in vitro* experiments were carried out with the three-electrode configuration in a 15-mL electrochemical glass cell (Bioanalytical Systems, InC). Cyclic voltammetry and Nyquist plot were measured in a K_3_Fe(CN)_6_/K_4_Fe(CN)_6_ (1:1, v/v, 5.0 mM) solution dissolved in PBS (50 mM) at pH 7.0. Electrochemical impedance spectroscopy (EIS) was characterized in PBS. Amperometry measurements were performed in PBS with constant stirring to characterize dopamine sensitivity. A constant potential of +0.4 V (vs. Ag/AgCl, 3M NaCl) was applied to the working electrode (rGO modified roughened Pt electrode) and the background current was stabilized before aliquots of dopamine were added to the electrochemical cell.

### COMSOL simulation for origami probe design

Simulation was performed using COMSOL Multiphysics® to evaluate different hinge designs in the origami neural probe. In the model, a pair of single- or double-layer shanks (60 µm × 400 µm) connected with thinned elastic hinges were modelled using polyimide material (Young’s Modulus: 8.5GPa). A periodic boundary condition was applied on the start and end surfaces (longitudinal direction) to simulate the full probe shank. A fixed boundary constraint was applied to one shank. A uniform pressure was applied onto the other shank to mimic the capillary force. A range of boundary load was simulated by parametric sweep. The displacement, bending angle, first principal strain and total elastic energy were simulated and evaluated.

### Acute implantation of MULTI and 3D origami neural probes

All animal handling and experimental procedures associated with and performed in this study followed the National Institutes of Health (NIH) animal use guidelines and were approved by the Institutional Animal Care & Use Committee (IACUC) at the University of Michigan (Approval Number PRO00009668). On the day of experimentation, the subject was transferred into an anesthetic induction chamber with a constant isoflurane flow at 5%. Immediately after, the subject was administered a urethane intraperitoneal injection with a dose of 1500 mg/kg. After confirming the anesthetic plane, the subject was administered with a subcutaneous atropine injection (0.05mg/kg) and secured into a stereotaxic frame with a heater set to 37℃. After removal and disinfection of the skin on the head, the skull was exposed and cleaned. The implant site was located and marked (Rat MULTI for temperature sensing: AP-3.24, ML-2.2 and DV-2.4; Rat MULTI for dopamine sensing: AP+1.92, ML-1.3 and DV-5.6; and Mouse for 3D µLED optogenetic stimulation and recording: AP-1.1mm, ML-1.0mm and DV-1.5mm). A stainless-steel ground screw was introduced into the occipital bone above the cerebellum and sealed in place with light cured acrylic (Unifast LC). A craniotomy was performed over the target area and the dura was removed. The device was mounted onto a shuttle and covered with PEG 12K to prevent it from buckling during insertion. The device was then slowly lowered until the target depth was reached. Once in place, the craniotomy was sealed with a two-part silicone and 15-20 minutes were given for the tissue to stabilize before recording. Implantation of the 3D recording and 3D µLED origami probes differ from the MULTI devices in the insertion shuttle, which was replaced by the fiber attached to the PCB. Recordings were acquired at 20,000 samples/s using Intan amplifier boards (multiple 32-channel or 128-channel headstages). For the 3D µLED origami probe experiment, each µLED was triggered 50 times using a source-meter (Keithley 2400) at 2Hz and 10% duty cycle (50ms pulse width). The stimulation sequence was controlled with a custom python script and synced via TTL to the recording board for further analysis.

### Chronic implantation and recording of stacked and double-sided neural probes

For the chronic implantations, the same anesthesia and surgical procedures as described above were performed first. A 3D printed base was cemented to the skull with light-cured acrylic (Unifast LC). Next, a ground screw was introduced, and the neural probe was inserted as described above. In addition, the rigid insertion shuttle was retrieved after PEG was fully dissolved. The craniotomy was sealed with soft two-part silicon and a layer of light cured acrylic. A faraday cage was built around the implant with metal posts and a copper mesh, the PCB was then secured to the cage’s frame, and ground and reference were soldered to the mesh along with the wire coming from the ground screw. Immediately after surgery and for 3 days after, the subjects were treated with Enrofloxacin (8 mg/kg) and Carprofen (5 mg/kg). Subjects were monitored daily until fully recovered. Recording sessions were acquired weekly at 20,000 samples/s in the home cage using dual Intan amplifier boards (32-channel headstages).

### Spike sorting, single unit activity, and sharp-wave-ripple analysis

Spike sorting was performed using Kilosort. Next, the auto-clustered spikes were further curated using Phy. During this process, electrical noise mis-determined as spike signals were removed. Spike clusters which were in fact the same unit were merged manually. The curated spike data was further processed using CellExplorer. With CellExplorer, electrophysiological information such as average peak-to-peak amplitudes, autocorrelation histograms, firing rates, and neural connectivity were extracted. Determining cell-body positions using trilateration was performed using CellExplorer from the recording data of MULTI and double-sided origami probe. For 3D recording, the trilateration was performed with CellExplorer using a 12-column staggered array for channel mapping. If signals of the same unit are shown on both the first and the last column, an additional trilateration is performed for these two columns. Lastly, the trilaterated locations were projected onto the circumference of a cylinder. Peristimulus time histograms (PSTH) were calculated for each neuron and for all the neurons per shank in 1ms bins. A 10ms window gaussian smoothing was applied to the resulting PSTH and normalized to the maximum value in the PSTH window (-10 - +70ms).

Local field potential extraction, bandpass filtering, and sharp-wave-ripple detection and analysis were performed using the open-source package from the Buzsáki lab at the New York University. Next, peak alignment was conducted using MATLAB. The minimum peak was set as the center of the extracted ripples for alignment. The signals with a 20-ms duration before and after the ripple center were captured. If less than 10% of randomly sampled ripples exhibited waveforms noticeably different from the average waveform and did not appear to reflect systematic bias, we did no further alterations to remove those signals.

## Acknowledgements

This research was supported by the National Institutes of Health (NIH), RF1NS133978 and RF1NS113283. We thank Dr. Eunah Ko for her expertise and suggestions with the flexible μLED probe design and fabrication. We thank Dr. John P. Seymour for his advice on flexible probe longevity improvement, Dr. Jinsang Kim for his advice and laboratory support in early phase 3D origami probe development, and Dr. Cynthia A Chestek for her advice on neural transmitter sensing. We also thank Dr. Sandrine Martin, Dr. Pilar Herrera-Fierro, Brian Armstrong, Shawn Wright and other staff members at the Lurie Nanofabrication Facility and Nancy Senabulya Muyanja at the Michigan Center for Materials Characterization for their technical advice and assistance in microfabrication and characterization.

## Supplementary information

**Table S1.** Chemical formula and concentration of Ni_80_Fe_20_ electroplating baths. A meshed Ni plate was used as an anode. DC current with a density at 10 mA/cm^2^ was applied during plating.

| *Chemical* | *Concentration* |
| --- | --- |
| Nickel(II) sulfate ($\text{NiSO}_4 \cdot 6\text{H}_2\text{O}$) | 200.0 g/L |
| Nickel(II) chloride ($\text{NiCl}_2 \cdot 6\text{H}_2\text{O}$) | 5.0 g/L |
| Iron(II) sulfate ($\text{FeSO}_4 \cdot 7\text{H}_2\text{O}$) | 8.0 g/L |
| Boric acid ($\text{H}_3\text{BO}_3$) | 25.0 g/L |
| Saccharin ($\text{C}_7\text{H}_5\text{NO}_3\text{S}$) | 3.0 g/L |

**Figure S1.**
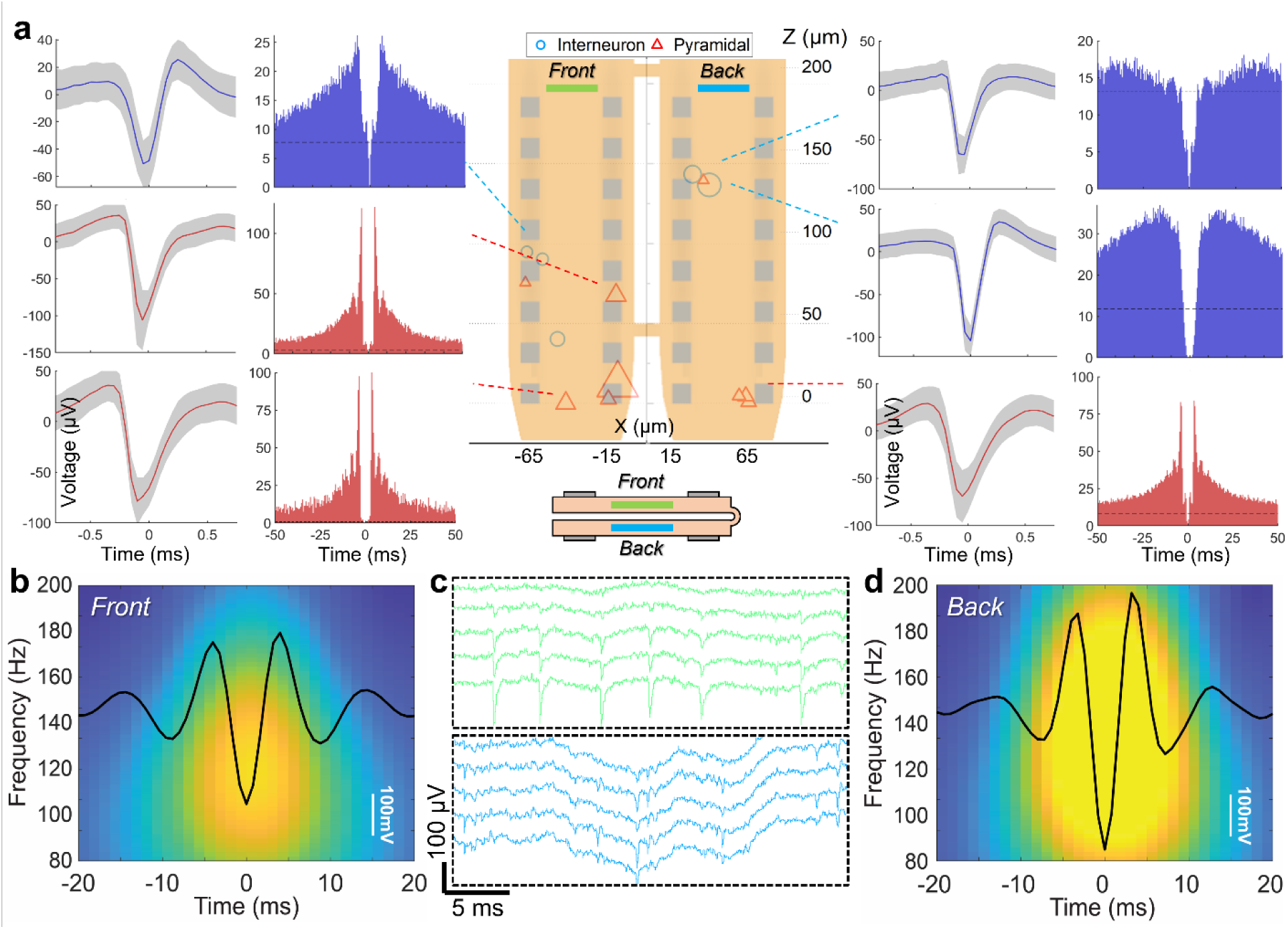
Neural recording with a double-sided probe (completely folded back-to-back) after 1-week implantation. **a.** Representative waveforms and auto-correlograms of the single units recorded on the front and back side. The symbol size represents the mean amplitude with an average value of 120.3 μV. **b.** and **d.** The average waveforms and power spectrogram of the sharp-wave-ripples recorded on the front and back side. **c.** Wide-band signals recorded from both sides with a duration of 30 ms.

**Figure S2.**
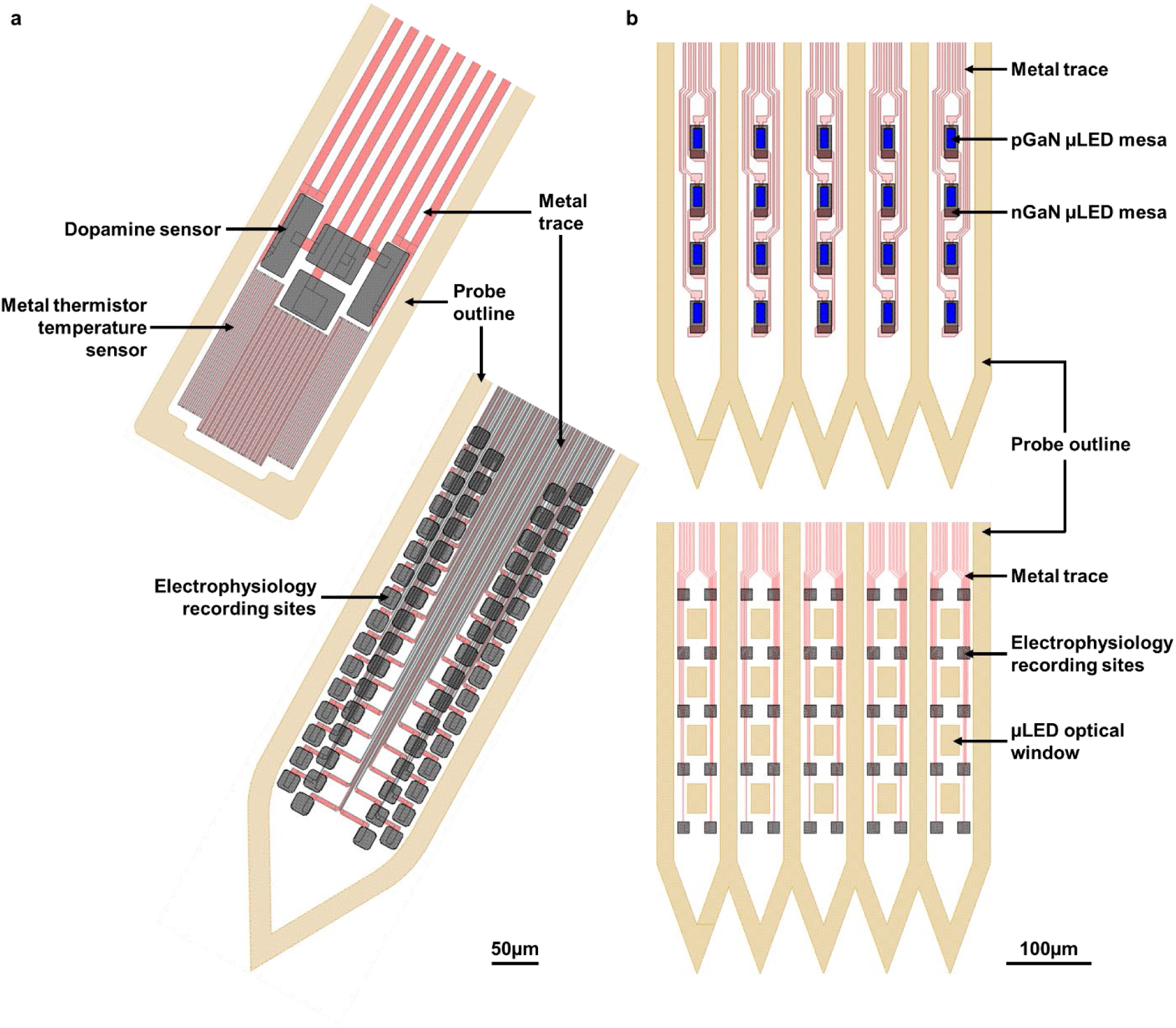
Layout of **a.** “temperature and dopamine sensing module (top)” and 64-channel “recording module (bottom),” and **b.** “3D origami µLED module (top)” and 50-channel “3D origami recording module (bottom).”

**Figure S3.**
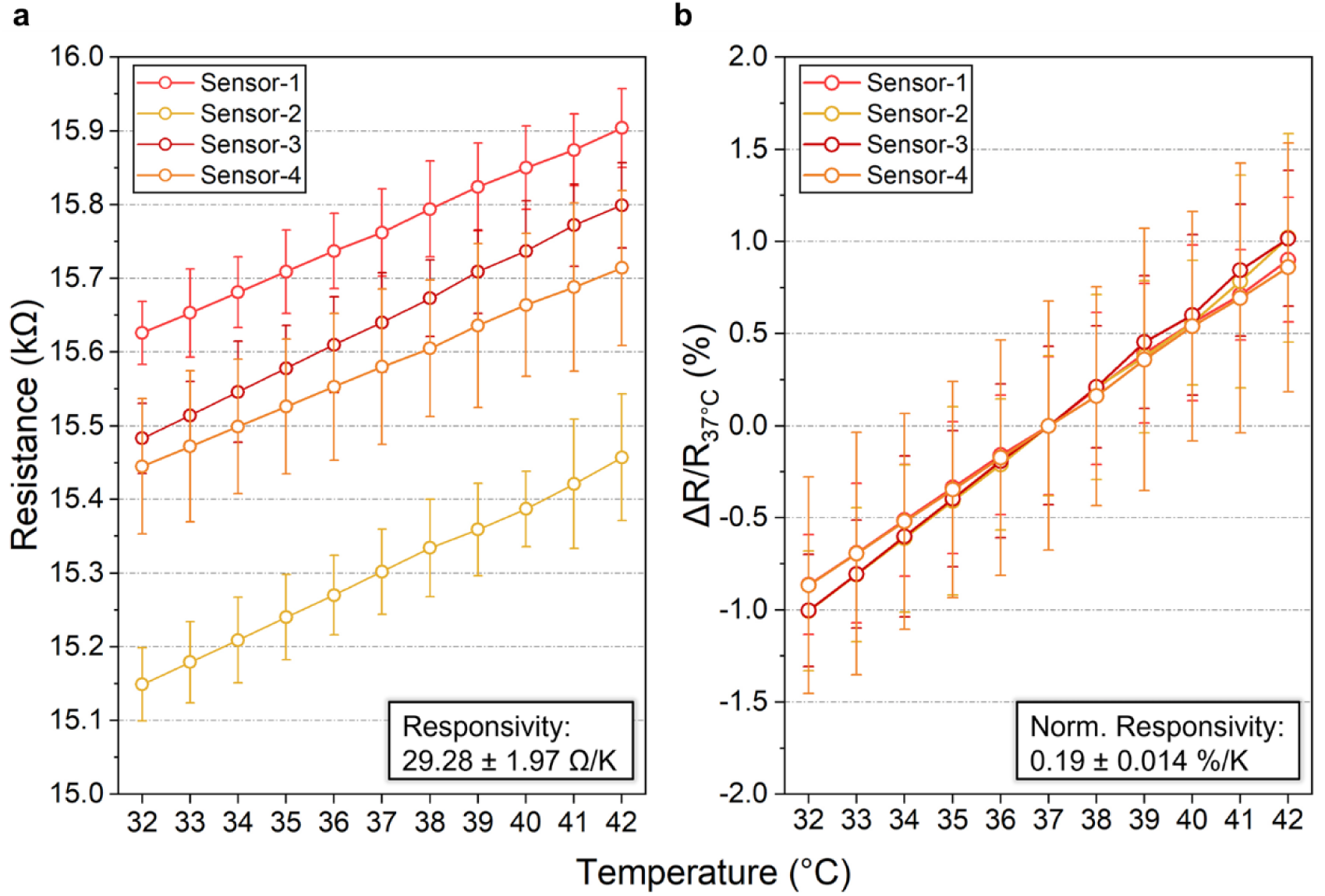
Characterization of the metal thermistor temperature sensor: **a.** responsivity and **b.** normalized responsivity.

**Figure S4.**
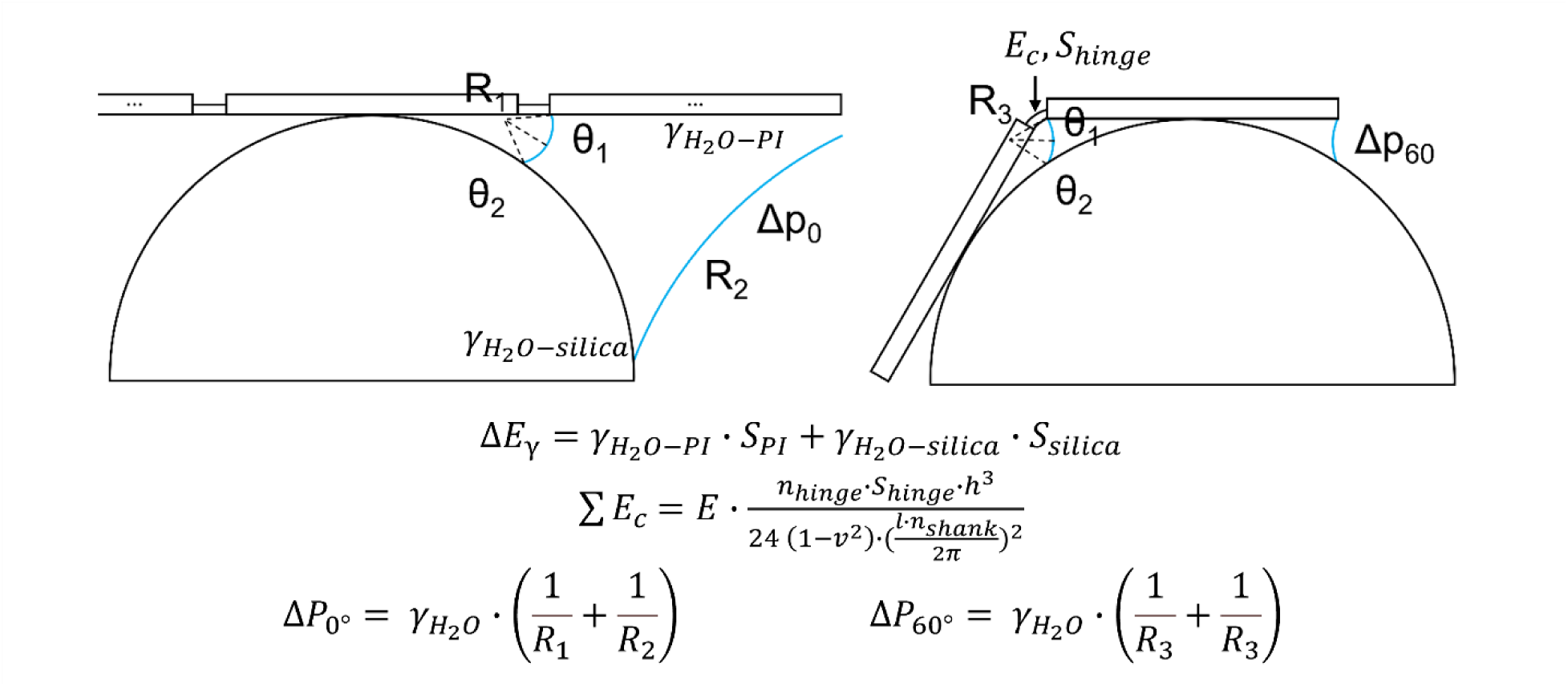
Estimation of the surface energy reduction, *E_γ_*, hinge elastic energy, *E_c_*, and capillary force induced pressure difference, *ΔP*, at the beginning (left) and ending (right) of the capillary-assisted wrapping process. *θ* is the contact angle of water on polyimide and silica; *S* is surface area of probe shank, optical fiber, and hinges; *γ* is surface energy density; *n* is the number of hinges and shanks; *l* and *h* are the length and thickness of the hinges, respectively; *R* is the radius of curvature; and *E* and *ν* are the Young’s modulus and Poisson ratio of polyimide, respectively. Based on the 6-shank design for assembly over a fiber core of 105-μm in diameter, the total estimated surface energy reduction is 13.9 nJ, and the capillary-induced pressure at the start and the end of wrapping are 9.1 kPa, and 15 kPa, respectively.

**Figure S5.**
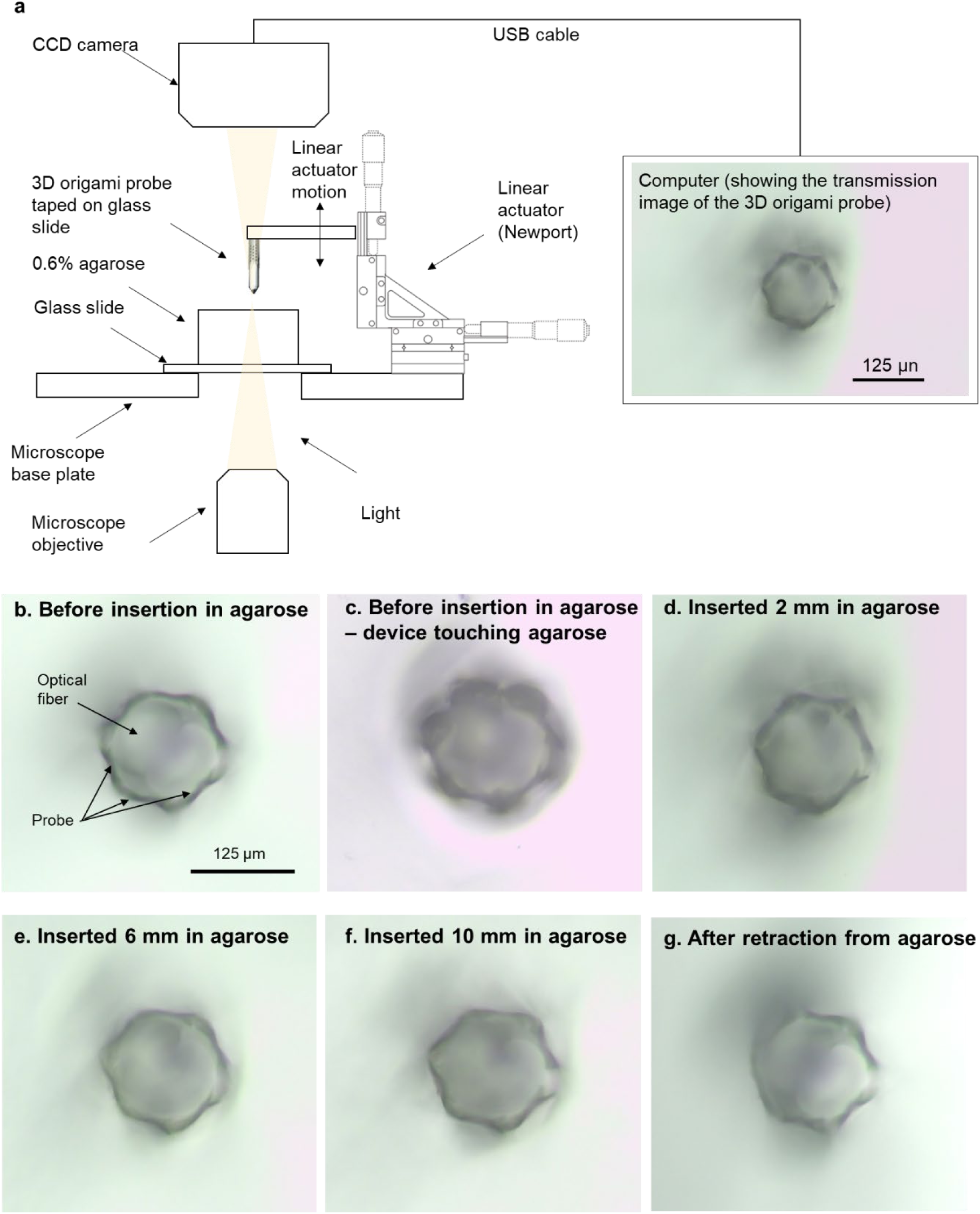
Insertion testing of the 3D origami probe. **a.** Setup schematic of experiment, **b-g.** Microscope photos of 3D probe at different insertion depths.

**Figure S6.**
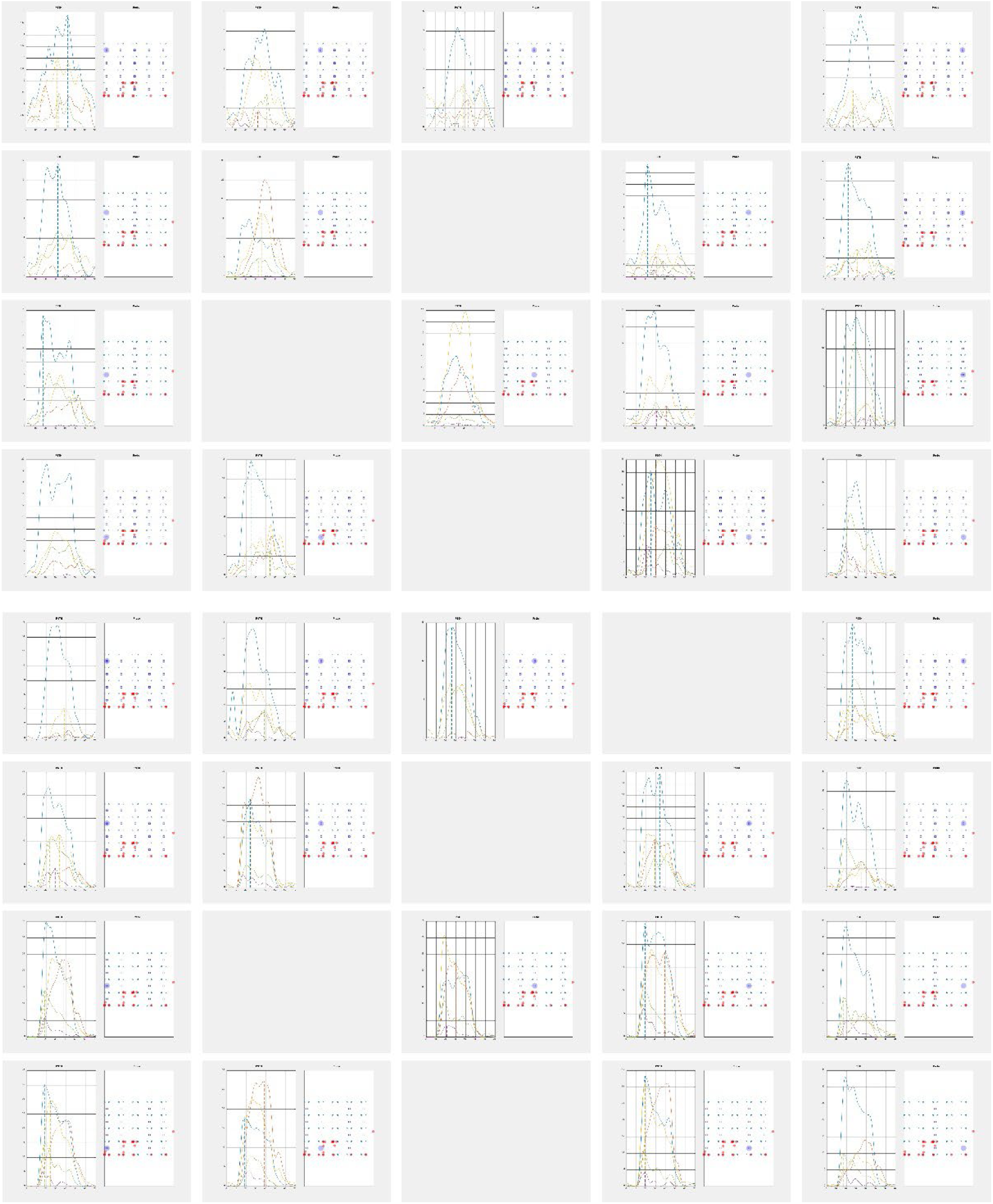
Per LED peristimulus time histograms. First 4 rows correspond to low power settings and 4 bottom rows to high power settings. PSTHs were calculated for each µLED by binning the APs from the corresponding neurons in each shank (in 1ms bins, APs within -2 to +3ms to the µLED onset were discarded), aligned to the µLED onset, and smoothened with a10ms gaussian. Shank color coding: S1=blue, S2=red, S3=yellow, S4=purple and S5=green. The location of all the recorded neurons and the triggered µLEDs are superimposed over the probe layout for each case.

**Figure S7.**
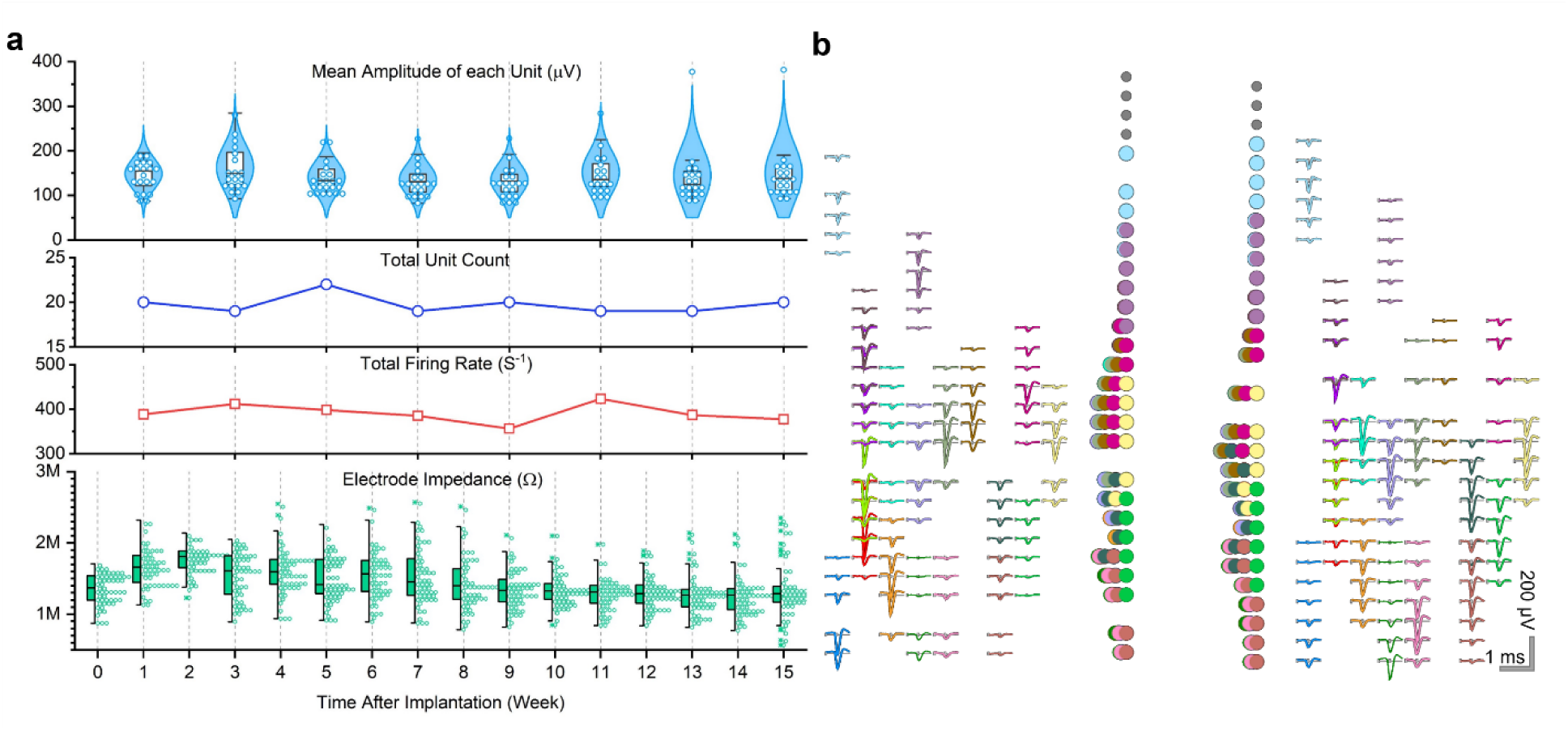
Chronic recording with a 64-channel stacked flexible probe. **a.** Box plot of the mean and maximum amplitude from each single unit. Total number of single units and total firing rates are shown for every other week and electrode impedance is shown for every week until week 15. **b.** Templates for all the single units recorded for every other week for a total of 15 weeks.

